# EMBER: Multi-label prediction of kinase-substrate phosphorylation events through deep learning

**DOI:** 10.1101/2020.02.04.934216

**Authors:** Kathryn E. Kirchoff, Shawn M. Gomez

## Abstract

Kinase-catalyzed phosphorylation of proteins forms the backbone of signal transduction within the cell, enabling the coordination of numerous processes such as the cell cycle, apoptosis, and differentiation. While on the order of 10^5^ phosphorylation events have been described, we know the specific kinase performing these functions for less than 5% of cases. The ability to predict which kinases initiate specific individual phosphorylation events has the potential to greatly enhance the design of downstream experimental studies, while simultaneously creating a preliminary map of the broader phosphorylation network that controls cellular signaling. To this end, we describe EMBER, a deep learning method that integrates kinase-phylogeny information and motif-dissimilarity information into a multi-label classification model for the prediction of kinase-motif phosphorylation events. Unlike previous deep learning methods that perform single-label classification, we restate the task of kinase-motif phosphorylation prediction as a multi-label problem, allowing us to train a single unified model rather than a separate model for each of the 134 kinase families. We utilize a Siamese network to generate novel vector representations, or an embedding, of motif sequences, and we compare our novel embedding to a previously proposed peptide embedding. Our motif vector representations are used, along with one-hot encoded motif sequences, as input to a classification network while also leveraging kinase phylogenetic relationships into our model via a kinase phylogeny-weighted loss function. Results suggest that this approach holds significant promise for improving our map of phosphorylation relations that underlie kinome signaling.

**Availability:** https://github.com/gomezlab/EMBER

## Introduction

Phosphorylation is the most abundant post-translational modification of protein structure, affecting from one to two-thirds of eukaryotic proteins. In humans, the number of kinases catalyzing this reaction hints at its importance, with kinases being one of the largest gene families with roughly 520 members distributed among 134 families (1–3). During phosphorylation, a kinase facilitates the addition of a phosphate group at serine, threonine, tyrosine, or histidine residues; though other sites exist. Phosphorylation of a substrate at any of these residues occurs within the context of specific consensus phosphorylation sequences, which we refer to here as “motifs”. Additional substrate binding sequences within the kinase or substrate, as well as protein scaffolds that facilitate structural orientation and downstream catalysis of the reaction, modify the efficacy of motif phosphorylation. Typically, the net effect of kinase phosphorylation is to switch the downstream target into an “on” or “off” state, enabling the transmission of information throughout the cell. Kinase activity touches nearly all aspects of cellular behavior, and the alteration of kinase behavior underlies many diseases while simultaneously establishing the basis for therapeutic interventions (4–11).

Although the importance of phosphorylation in cell information processing and its dysregulation as a driver of disease is well-recognized, the map of kinase-motif phosphorylation interactions is mostly unknown. So, while upwards of 100,000 motifs are known to be phosphorylated, less than 5% of these have an associated kinase identified as the catalyzing agent (12). This knowledge gap provides a considerable impetus for the development of methods aimed at predicting kinase-motif phosphorylation events that, at a minimum, could help focus experimental efforts.

As a result, a number of computational tools have been developed, spanning a myriad of methodological approaches including random forests (13), support vector machines (14), logistic regression (15), and Bayesian decision theory (16). Advances in deep learning have similarly spawned new approaches, with two methods recently described. The first, MusiteDeep, utilizes a convolutional neural network (CNN) with attention to generate single predictions (17). The second deep learning method, DeepPhos, exploits densely connected CNN (DC-CNN) blocks for its predictions (18). Both of these approaches train individual models for each kinase family, requiring a separate model for each of the 134 kinase families. In addition to the practical challenge of training many individual models, a further disadvantage of these two deep learning approaches is the potential lost opportunity gains from transfer learning, as models trained independently do not directly incorporate knowledge of motif phosphorylation by kinases from different kinase families.

Here, we describe, EMBER (<u>E</u>mbedding-based <u>m</u>ulti-la<u>b</u>el pr<u>e</u>diction of phospho<u>r</u>ylation events), a deep learning approach for predicting *multi-label* kinase-motif phosphorylation relationships. In our approach, we utilize a Siamese neural network, modified for our multi-label prediction task, to generate a high-dimensional embedding of motif vectors. We further utilize one-hot encoded motif sequences. These two representations are leveraged together as a dual input into our classifier, improving learning and prediction. We also find that our Siamese embedding generally outperforms a previously proposed protein embedding, ProtVec, which is trained on significantly more data (19). We further integrate information regarding evolutionary relationships between kinases into our classification network loss function, informing predictions in light of the sparsity associated with these data, and we find that this information improves prediction accuracy. As EMBER utilizes transfer learning across families, we expect that model accuracy will improve more so than other deep learning approaches as more data describing kinase-substrate relationships are collected. Together, these results suggest that EMBER holds significant promise for improving our map of phosphorylation relationships that underlie the kinome and broader cellular signaling.

## Methods

### Kinase-motif interaction data

As documented kinase-motif interactions are sparse in relation to the total number of known phosphorylation events, we attempted to maximize the number of examples of such interactions for training. To do this, we integrated multiple datasets describing kinase-motif relationships across multiple vertebrate species. Our data was sourced from PhosphoSitePlus, PhosphoNetworks, and Phospho.ELM, all of which are collections of annotated and experimentally verified kinase-motif relationships (20–22). From these data sources, non-redundant kinase-motif relationships were extracted and integrated into a single set of interactions. We used the standard single-letter amino acid code for representation of amino acids, with an additional ’X’ symbol to represent an ambiguous amino acid. We defined our motifs as peptides composed of a central phosphorylatable amino acid — either serine (S), threonine (T), or tyrosine (Y) — flanked by seven amino acids on either side. Therefore, each motif is a 15-amino acid peptide or “15-mer”. As a phosphorylatable amino acid may not have seven flanking amino acids to either side if it is located near the end of a substrate sequence, we used ‘-’ to represent the absence of an amino acid in order to maintain a consistent motif length of 15 amino acids across all instances.

Deep learning models are known to generally require large amounts of examples per class in order to achieve adequate performance. Our original dataset was considerably imbalanced in that all positive labels (verified kinase-motif interactions) had a very low positive-to-negative label ratio. For example, the TLK kinase family only has nine positive labels (verified TLK-motif interactions) and more than 10,000 negative labels (lack of evidence for a TLK-motif interaction). To maximize our ability to learn from our data, we utilized only kinases that had a relatively large number of experimentally validated motif interactions, reducing the number of kinase-motif relationships to be used as input for our model. This filtering also served to considerably mitigate the label imbalances in our data. From the 7531 remaining motifs, we set aside 853 motifs for the independent test set, leaving 6678 for the training set. Then we removed any sequences from the training set that met a 60% similarity threshold with any sequence in the test set, based on Hamming distance scores. This process removed 229 motifs from the training set. Kinase labels were then grouped into respective kinase families contingent on data collected from the RegPhos (1) database, resulting in eight kinase families. Our resulting data set is comprised of 7302 phosphorylatable motifs and their reaction-associated kinase families (Table 1). Furthermore, our data are multi-label in that a single motif may be phosphorylated by multiple kinases, including those from other families, resulting in a data point with potentially multiple positive labels.

**Table 1.** Summary of our kinase-motif phosphorylation dataset. Shown are the number of kinases per family along with the number of motifs phosphorylated by each kinase family in the training and test sets.

| Family | Kinases | Training motifs | Testing motifs |
| --- | --- | --- | --- |
| Akt | 3 | 382 | 63 |
| CDK | 21 | 752 | 116 |
| CK2 | 2 | 775 | 97 |
| MAPK | 14 | 1275 | 187 |
| PIKK | 7 | 497 | 63 |
| PKA | 5 | 1235 | 231 |
| PKC | 10 | 1497 | 251 |
| Src | 11 | 869 | 99 |

### Motif embeddings

#### ProtVec embedding

We chose to investigate two methods to achieve our motif embedding. First, we considered ProtVec, a learned embedding of amino acids, originally intended for protein function classification (19). ProtVec is the result of a Word2Vec algorithm trained on a corpus of 546,790 sequences obtained from Swiss-Prot, which were broken up into 3 amino acid-long subsequences, or “3-grams”. As a result of this approach, ProtVec provides a 100-dimensional distributed representation, analogous to a natural language “word embedding”, that establishes coordinates for each possible amino acid 3-gram. This results in a 9048 × 100 matrix of coordinates, one 100-dimensional coordinate for each 3-gram. In a preliminary investigation, we found that averaging the ProtVec coordinates resulted in a higher-quality embedding compared to the original ProtVec coordinates. Comparisons between the two embeddings are provided in Supplemental Material. We averaged the embedding coordinates, per amino acid, in the following fashion:

We define *T* = [`AAA, ALA, LAA, …, unknown`], the vector of 9048 amino acid 3-grams provided by the authors of ProtVec. We also define *A* = [`A, L, S`, …, -], the alphabet comprising the 22 amino acid symbols. We equate “-” to the “unknown” character defined by ProtVec. Then, we compute the matrix of averaged ProtVec coordinates, *C*^(*avg*)^, which will be 22 × 100 dimensions:

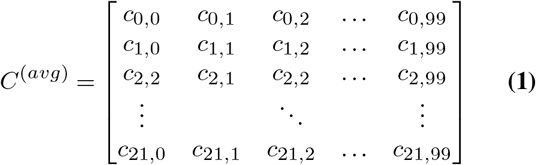

We solve for each element of *C*^(*avg*)^ based on the values of *C*^(*raw*)^, the original (9048 × 100) ProtVec matrix:

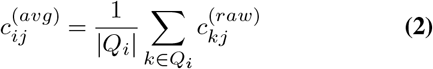

where 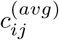 belongs to *C*^(*avg*)^, 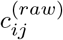 belongs to *C*^(*raw*)^, and

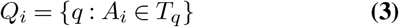

Note that the original ProtVec matrix was 9048 × 100 dimensions, thus each *j* corresponds to the index of one of the 100 original ProtVec dimensions along the second tensor dimension.

#### Siamese embedding

We aimed to produce a final model, composed of an embedding technique and a classification method, that was specific to our motif dataset. To this end, we implemented a Siamese network to provide a novel learned representation of our motifs (Figure 1). The Siamese network is composed of two identical “twin” networks, deemed as such due to their identical hyperparameters as well as their identical learned weights and biases (23). During training, each twin network receives a separate motif sequence that is represented as a one-hot encoding, denoted either as *a* or *b* in Figure 1. Motifs are processed through the network until reaching the final fully-connected layers, *h_a_* and *h_b_*, which provide the resultant embeddings for the original motif sequences. Next, the layers are joined by calculating the pair-wise Euclidean distance, *D_w_*, between *h_a_* and *h_b_*. *D_w_* can be interpreted as the overall dissimilarity between the original motif sequences, *a* and *b*. The loss function operates on the final layer, striving to embed relatively more similar data points closer to each other, and relatively more different data points farther away from each other. In this way, the network amplifies the similarities and differences between motifs, and it translates such relationships into a semantically meaningful vector representation for each motif in the embedding space. We utilized a contrastive loss as described in Hadsell et al. (24), but we sought to modify the function to account for the multi-label aspect of our task. The canonical Siamese loss between a pair of samples, *a* and *b*, is defined as

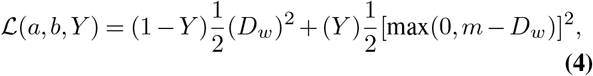

where *D_w_* is the Euclidean distance between the outputs of the embedding layer, *m* is the margin which is a hyperparameter defined prior to training, and *Y* ∈ {0, 1}. The value of *Y* is determined by the label of each data point in the pair. If a pair of samples has *identical* labels, they are declared “same” (*Y* = 0). Conversely, if a pair of samples has *different* labels, they are declared “different” (*Y* = 1). This definition relies on the assumption that each sample may only have one true label. To adapt the original Siamese loss to account for the multi-label aspect of our task, we replaced the discrete variable *Y* with a continuous variable, namely, the Jaccard distance between kinase-label set pairs. Thus, our modified loss function is defined as

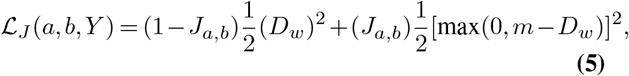

where *J_a,b_* is shorthand for *J* (*K_a_, K_b_*), which is the Jaccard distance between the kinase-label set *K_a_* and the kinase-label set *K_b_*, associated with motif sample *a* and motif sample *b*, respectively. Formally,

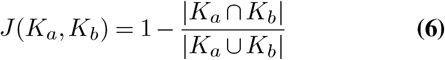

and consequently,

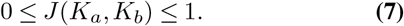

**Fig. 1.**
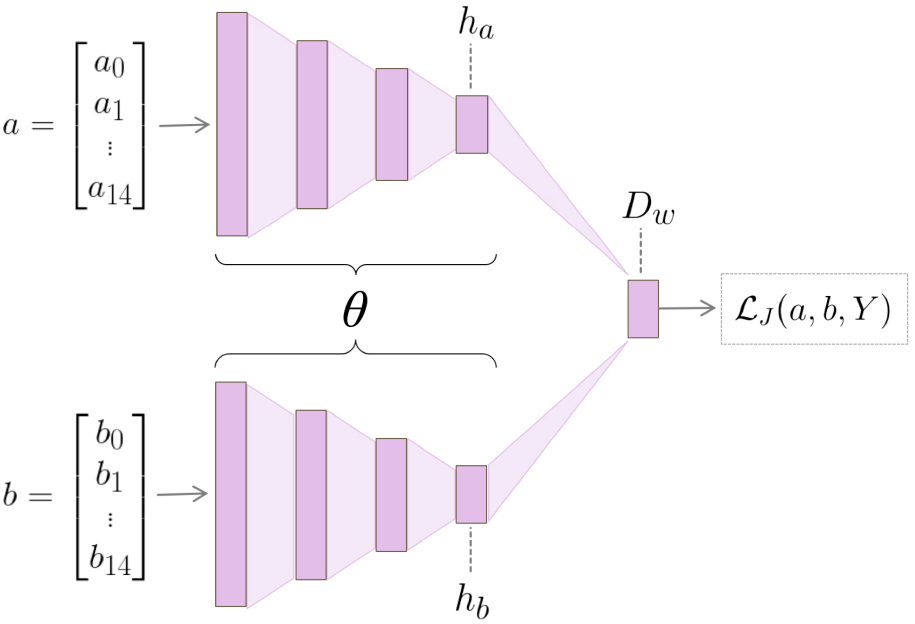
Siamese network architecture, composed of twin convolutional neural networks (CNNs). The twin networks are joined at the final layer. *a* and *b* represent a pair of motifs from the training set, while *h_a_* and *h_b_* represent the respective hidden layers output by either CNN. The difference between the hidden layers is calculated to obtain the distance layer, *D_w_*. *D_w_* is input into the loss along with *Y*, a variable indicating the dissimilarity, regarding kinase interactions, between *a* and *b*. After training is complete, the “twin” architecture is no longer necessary; each motif is input into a single twin and the output of the embedding layer gives the resultant representation of the given motif.

In this way we have defined a continuous metric by which to compare a pair of motifs, rather than the usual “0” or “1” distinction.

The Siamese network was trained for 10,000 iterations on the training set, precluding the data points in the independent test set. When composing a mini-batch, we alternated between “similar” and “dissimilar” motif pairs during training. Similar pairs were defined as motifs whose *J* (*K_a_, K_b_*) > 0.5, and dissimilar pairs were defined as motifs whose *J* (*K_a_, K_b_*) ≤ 0.5. After training, we must produce the final embedding space to be used in training of our subsequent classification network. To obtain the final embedding, we input each motif into a single arbitrary twin of the original network (because both twins learn the same weights and biases), producing a high-dimensional (100-dimensional) vector representation of the original motif sequence. The resultant motif embedding effected by the single Siamese twin is further discussed in the Results section. We used k-nearest neighbors (*k*-NN) classification on each family to quantitatively compare the predictive capabilities of ProtVec and Siamese embeddings in the coordinate-only space. For our *k*-NN computation, we used a *k* of 85.

### Predictive model framework

#### EMBER architecture

An overview of the architecture of EMBER is shown in Figure 2. EMBER takes as input raw motif sequences and the coordinates of each respective motif in the embedding space. We use one-hot encoded motifs as the second type of input into our model. Each motif sequence is represented by a 15 × 22 matrix. In addition, we utilize the embedding provided by our Siamese network, which creates a latent space of dimensions *m* × 90 where *m* is the number of motifs.

**Fig. 2.**
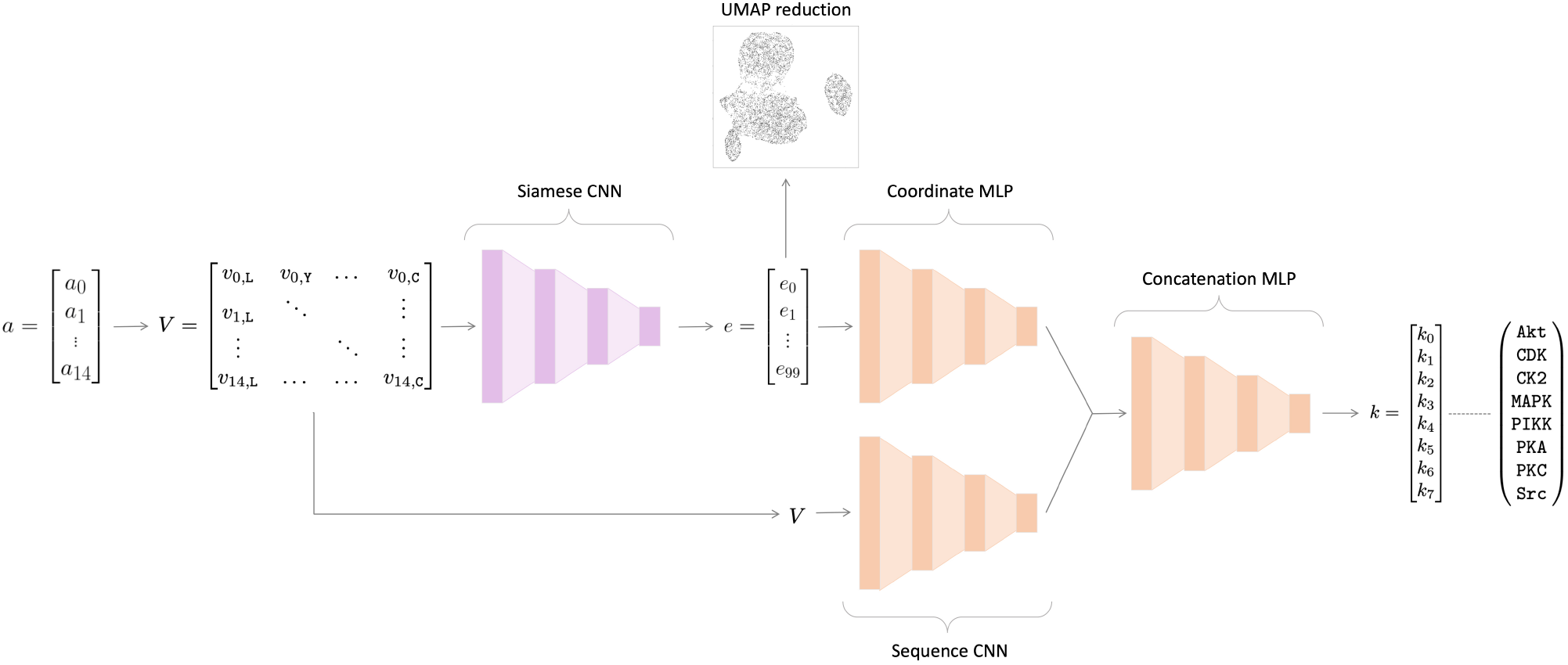
EMBER model architecture. Here, the previously-trained Siamese network is colored pink, and the classifier architecture is colored orange. The 15 amino acid-length motif, *a*, is converted into a one-hot encoded matrix, *V*. The one-hot encoded matrix is then fed into a single twin from the Siamese network. The 100-d embedding, *e*, is output by the Siamese network. Here, we reduce *e* to a 2-dimensional space for illustrative purposes using UMAP. Then, *e* is fed into a multilayer perceptron (MLP) alongside *V*, which is fed into a convolutional neural network (CNN). Then, the last layers of the separate networks are concatenated, followed by a series of fully-connected layers. The final output is a vector, *k*, of length eight, where each value corresponds to the probability that the motif *a* was phosphorylated by one of the kinase families indicated in *k*.

The inputs into our classifier network, one-hot sequences and embeddings, are fed through a convolutional neural network (CNN) and a multilayer perceptron (MLP), respectively. The outputs of the two networks are then concatenated, and the concatenated layer is fed through a series of fully-connected layers (a MLP), followed by a sigmoid activation function. We performed 5-fold cross validation to assess the accuracy of our model when trained on different training-validation folds. We averaged the performance on the independent test set across the five folds to compute our final performance on the classification task.

#### Evaluation metrics

In order to quantify the performance of our models, we computed the area under the receiver operating characteristic curve (AUROC) and the area under the precision-recall curve (AUPRC). These metrics were evaluated per kinase family. We also show the micro-average and macro-average for both AUROC and AUPRC. We define Λ = {*λ_j_* : *j* = 0, …, *q*} as the set of all labels. The micro-average, *E_micro_*, aggregates the label-wise contributions of each class:

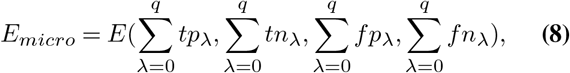

where *E* is an evaluation metric, in our case, either AUROC or AUPRC. Alternatively, the macro-average, *E_macro_*, takes into account the score for each respective class and averages those scores together, thus treating all classes equally:

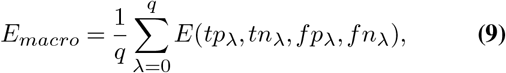

where *E* is once again an evaluation metric, in our case, either AUROC or AUPRC. Both the *E_macro_* and the *E_micro_* are calculated based on *tp_λ_*, *tn_λ_*, *fp_λ_*, and *fn_λ_*, which are, respectively, the number of true positives, the number of true negatives, the number of false positives, and the number of false negatives of label *λ*.

### Kinase phylogenetic distances

We sought to leverage the phylogenetic relationships between kinases to improve predictions of kinase-motif interactions. Specifically, we considered the dissimilarity of a pair of kinase families in conjunction with the dissimilarity of the two respective groups of motifs that either kinase family phosphorylates (i.e., “kinase-family dissimilarity” vs. “motif-group dissimilarity”). Note that the terms “distance” and “dissimilarity” are interchangeable. As the phylogenetic distances given by Manning et al. (2) do not provide distances between typical and atypical kinase families, we established a proxy phylogenetic distance that allows us to define distances between these two families. We define this proxy phylogenetic distance through the Levenshtein edit distance, *Lev*(*k_a_, k_b_*), between kinase-domain sequences. Kinase-domain sequences are the specific subsequences of kinases that are directly involved in phosphorylation. These kinase-domain sequences were obtained from an online source provided by Manning et al. (2). Distances between kinase domain sequences was calculated by performing local alignment, utilizing the BLOSUM62 substitution matrix to weight indels and substitutions. To calculate overall kinase-family dissimilarity, we took the average of the Levenshtein edit distances between each kinase domain pair, per family,

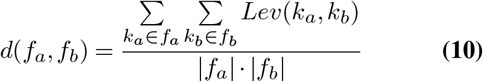

where *d*(*f_a_, f_b_*) is the dissimilarity metric (distance) between kinase family *a* and kinase family *b*. *k_a_* is the kinase-domain sequence of a kinase belonging to family *a*, *k_b_* is the kinase-domain sequence of a kinase belonging to family *b*, and the Levenshtein distance between kinase domain *k_a_* and kinase domain *k_b_* is determined by *Lev*(*k_a_, k_b_*). This formula was applied per kinase family pair and stored in an *a × b* kinase-family dissimilarity matrix. We will refer to this proxy metric for evolutionary dissimilarity between kinase families as the “phylogenetic distance” between kinase families.

#### Kinase-family dissimilarity vs. motif-group dissimilarity

For our (kinase-family dissimilarity)-(motif-group dissimilarity) correlation, we defined motif-group dissimilarity in the same manner as kinase-family dissimilarity, finding the Levenshtein distance between motifs based on local alignment using BLOSUM62. Then, we sought to find the correlation between kinase-family dissimilarity and motif-group dissimilarity. Therefore, calculation of motif-group dissimilarity, per kinase family pair, was defined identically as in Equation 10, but based on the motifs specific to each kinase family, resulting in an *a × b* motif-group dissimilarity matrix.

#### Kinase phylogenetic loss

To leverage evolutionary relationships between kinase families into our predictions, we weighted the original binary cross entropy (BCE) loss by a kinase phylogenetic metric. Specifically, our weighted BCE loss per minibatch is defined as:

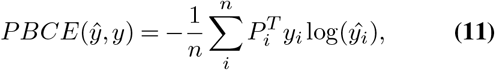

where *n* is the size of the mini batch, *y_i_* is the one-hot actual label vector for sample *i*, 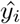 is the predicted label vector for sample *i*, and *P_i_* is the phylogenetic weight vector for sample *i* given by

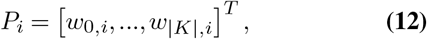

with *w_k,i_* being the average phylogenetic weight scalar of label *k* for sample *i*:

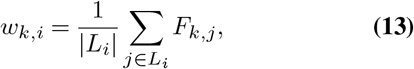

and *F_k,j_* is the vector of family weights of label *k*. Finally, *L_i_* is the set of indices corresponding to positive labels for sample *i*

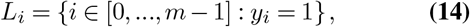

where *m* is the length of the one-hot true label vector for sample *i*.

## Results

### Correlation between kinase phylogenetic dissimilarity and phosphorylated motif dissimilarity

We sought to illuminate the relationship between kinase-family dissimilarity and phosphorylated motif-group dissimilarity described in the Methods section. That is, we wanted to determine if “similar” kinases tend to phosphorylate “similar” motifs based on some quantitative metric. To this end, we calculated the correlation between average kinase-family dissimilarities and motif-group dissimilarities based on normalized pairwise alignment scores. From this, we found a Pearson correlation of 0.667, indicating a moderate positive relationship between kinase dissimilarity and that of their respective phosphorylated motifs. While moderate, this correlation between kinase dissimilarity and motif dissimilarity suggests a potential signal in the phylogenetic relationships that could be leveraged to improve predictions.

Using our normalized distances as a proxy for phylogenetic distance (see Methods), the dissimilarity between kinases is displayed as a heatmap in Figure 3. The Akt and PKC family have the greatest similarity (lowest dissimilarity) of all pairwise comparisons, with PKA-Akt and MAPK-CDK following as the next most similar family pairs. Together, these results provide motivation to incorporate both motif dissimilarity and kinase relatedness into the predictive model, as achieved through our custom phylogenetic loss function described in Methods. The effects of this approach are described later in Results.

**Fig. 3.**
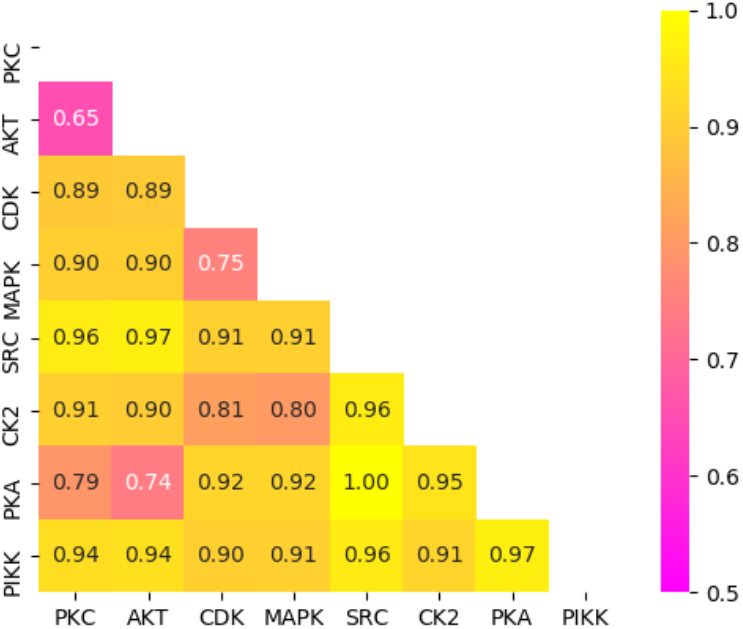
Heatmap matrix depicting pairwise kinase-domain distances. Levenshtein distances were normalized, with the yellow end of the color bar representing far distances (less similar) and the pink end representing close distances (more similar).

### Motif embedding via Siamese network

We sought to develop a novel learned representation of motifs using a Siamese neural network. Siamese networks were first introduced in the early 1990s as a method to solve signature verification, posed as an image-to-image matching problem (23). Siamese networks perform metric learning by exploiting the dissimilarity between a pair of data points. Training a Siamese network effects a function with the goal of producing a meaningful embedding, capturing semantic similarity in the form of a distance metric. We hypothesized that incorporating high-dimensional vector representations of motifs (i.e., an embedding) into the input of a classification network would provide more predictive power than methods that do not utilize such information. In our Siamese model, we opted to use convolutional layers as described in Methods. We performed *k*-NN on both the ProtVec and Siamese embeddings of motifs and found that the Siamese embedding produced better predictions, on average, than the ProtVec embedding (see Table 2). More specifically, the Siamese embedding resulted in a macro-average AUROC of 0.903 compared to ProtVec’s 0.898 and a micro-average AUROC of 0.924 compared to ProtVec’s 0.902. Likewise, the Siamese embedding had better AUPRC, with a macro-average AUPRC of 0.692 compared to ProtVec’s 0.670 and a micro-average AUPRC of 0.747 compared to ProtVec’s 0.643. Furthermore, we calculated the silhouette scores of both embeddings and found our Siamese embedding to have a significantly better mean silhouette score of 0.114 compared to ProtVec’s 0.005.

**Table 2.** Area under the receiver operating characteristic curve (AUROC) and area under the precision recall curve (AUPRC) scores on independent test set prediction, given by *k* -NN performed on the ProtVec and Siamese embedding.

| Family | Precision |  | Recall |  |
| --- | --- | --- | --- | --- |
|  | ProtVec | Siamese | ProtVec | Siamese |
| Akt | 0.908 | 0.897 | 0.462 | 0.513 |
| CDK | 0.889 | 0.892 | 0.511 | 0.538 |
| CK2 | 0.906 | 0.893 | 0.665 | 0.714 |
| MAPK | 0.907 | 0.908 | 0.739 | 0.720 |
| PIKK | 0.845 | 0.900 | 0.579 | 0.663 |
| PKA | 0.865 | 0.852 | 0.716 | 0.659 |
| PKC | 0.865 | 0.885 | 0.697 | 0.741 |
| Src | 0.998 | 0.995 | 0.993 | 0.991 |
| macro-average | 0.898 | **0.903** | 0.670 | **0.692** |
| micro-average | 0.902 | **0.924** | 0.643 | **0.747** |

We performed dimensionality reduction for visualization of the Siamese embeddings using uniform manifold approximation and projection (UMAP) (25). For our UMAP implementation, we used 200 neighbors, a minimum distance of 0.1, and Euclidean distance for our metric. The resulting 2-dimensional UMAP motif embeddings derived from the Siamese network are shown in Figure 4. As can be seen, the motifs phosphorylated by a given kinase family have a distinctive distribution in the embedding space, with some distributions being highly unique, and with some significant overlap between certain families. More specifically, our Siamese embedding shows that motifs phosphorylated by either PKC, PKA, or Akt appear to occupy a similar latent space. Similarly, motifs phosphorylated by either CDK or MAPK also occupy a similar space. These observations mirror the phylogenetic relationships shown in Figure 3, where the MAPK and CDK families have a relatively short mean evolutionary distance between them, and the PKC-PKA distance, even shorter still.

**Fig. 4.**
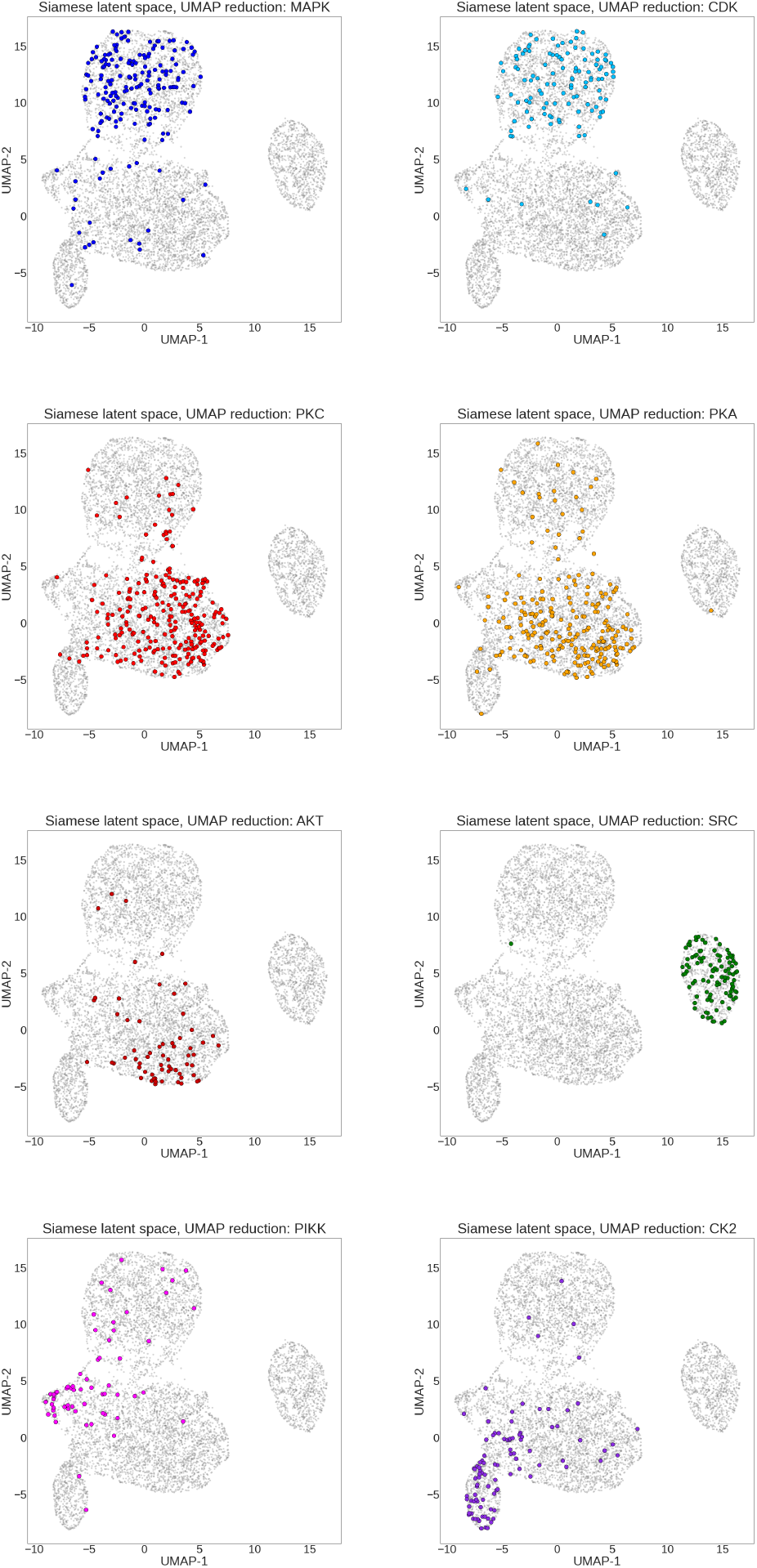
Siamese embedding of motifs. Each point represents one of the 7302 motifs, and each panel displays kinase family-specific phosphorylation patterns. Each colored point corresponds to a motif in the test set phosphorylated by a member of the specified kinase family. Highlighted points are slightly enlarged in size to enhance readability.

In addition to these overlapping families, we also observe that Src-phosphorylated motifs form a distinct cluster. This is likely driven by the fact that Src is the only tyrosine kinase family among the eight kinase families we investigated, with its motifs invariably having a tyrosine (Y) at the eighth position in the 15-amino acid sequence, compared to the other 7 families whose motifs have either a serine (S) or a in this position. This effects a significant sequence discrepancy between Src-phosphorylated motifs and remaining motifs. The fact that Src-phosphorylated motifs cluster so precisely serves as a sanity check that our Siamese embedding is capturing sequence (dis)similarity information despite being trained through comparison of kinase-motif phosphorylation events in lieu of motif sequence comparisons. We note that the embedding produced by our Siamese network is quite qualitatively similar to the ProtVec embedding in terms of these kinase-label clusters indicated in the UMAP projections. The UMAP projections of the ProtVec embeddings are included in Supplementary Material.

### Prediction of phosphorylation events

Following training of EMBER on both motif sequences and motif vector representations as input, we conducted an ablation test in which we removed the motif vector representation (or coordinate) input along with its respective MLP; this was achieved by applying a dropout rate of 1.00 on the final layer of the coordinate-associated MLP. This ablation test allowed us to observe how our novel motif sequence-coordinate model compares to a canonical deep learning model whose input consists solely of one-hot encoded motif sequences (such as in the methods utilized by Wang et al. (17) and Luo et al. (18)). We also compared EMBER trained with the standard BCE loss to EMBER trained with our kinase phylogenetic loss. All predictive models, as described in Table 3, were trained on identical training-validation splits and evaluated on the same independent test set.

**Table 3.**
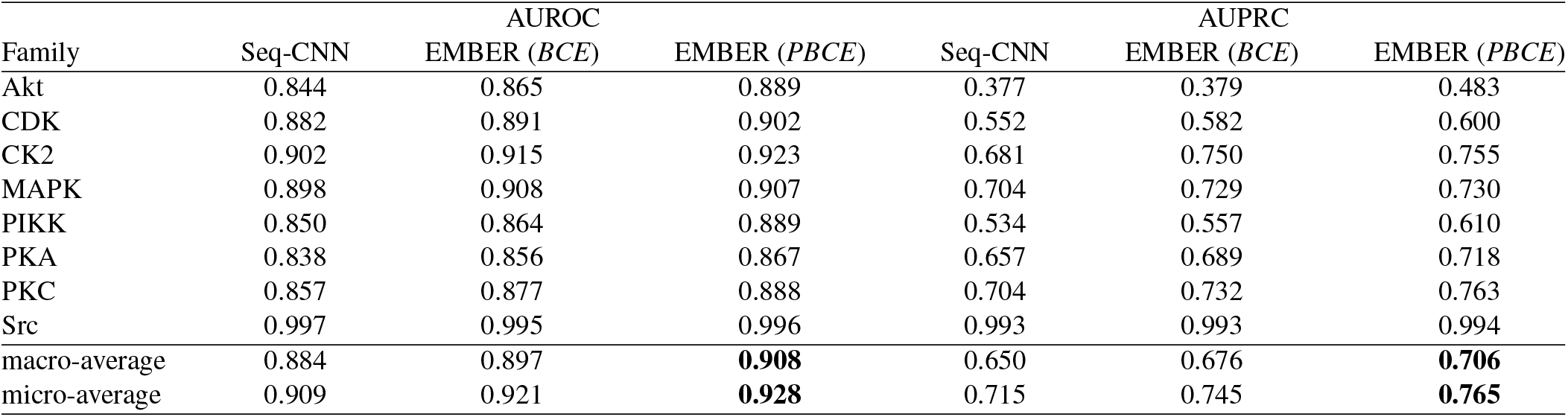
AUROC and AUPRC results achieved on the independent test set across deep learning classification models. The AUROC and AUPRC are presented per kinase family for each model. From left to right, we include results for the ablated sequence-only CNN, EMBER trained using a canonical BCE loss, and EMBER trained using the kinase phylogeny-weighted loss as described in Methods.

| Family | AUROC |  |  | AUPRC |  |  |
| --- | --- | --- | --- | --- | --- | --- |
|  | Seq-CNN | EMBER (*BCE*) | EMBER (*PBCE*) | Seq-CNN | EMBER (*BCE*) | EMBER (*PBCE*) |
| Akt | 0.844 | 0.865 | 0.889 | 0.377 | 0.379 | 0.483 |
| CDK | 0.882 | 0.891 | 0.902 | 0.552 | 0.582 | 0.600 |
| CK2 | 0.902 | 0.915 | 0.923 | 0.681 | 0.750 | 0.755 |
| MAPK | 0.898 | 0.908 | 0.907 | 0.704 | 0.729 | 0.730 |
| PIKK | 0.850 | 0.864 | 0.889 | 0.534 | 0.557 | 0.610 |
| PKA | 0.838 | 0.856 | 0.867 | 0.657 | 0.689 | 0.718 |
| PKC | 0.857 | 0.877 | 0.888 | 0.704 | 0.732 | 0.763 |
| Src | 0.997 | 0.995 | 0.996 | 0.993 | 0.993 | 0.994 |
| macro-average | 0.884 | 0.897 | **0.908** | 0.650 | 0.676 | **0.706** |
| micro-average | 0.909 | 0.921 | **0.928** | 0.715 | 0.745 | **0.765** |

Comparisons between the predictive capability of the models described here are quantified by AUROC and AUPRC, and these metrics are presented for each of the three models in Table 3. As indicated by Table 3, EMBER, utilizing both sequence and coordinate information, outperforms the canonical sequence model in both AUROC and AUPRC. In addition, integration of phylogenetic information into the loss provides a generally small but consistent additional boost in performance, showing the best overall results out of the three models for AUROC and AUPRC. Individual performance metric curves for each kinase label, produced by EMBER trained via the phylogenetic loss, are shown in Figure 6.

A confusion matrix providing greater detail and illustrating the relative effectiveness of our model for prediction of different kinase families is shown in Figure 5. In order to compute the confusion matrix, we set a prediction threshold of 0.5, declaring any prediction above 0.5 as “positive” and any prediction equal to or less than 0.5 as “negative”. As indicated by the confusion matrix, the model often confounds motifs that are phosphorylated by closely related kinase families, for example, MAPK and CDK. This is presumably due to the close phylogenetic relationship between MAPK and CDK, as indicated by their relatively low phylogenetic distance of 0.75 (Figure 3). Furthermore, our Siamese network embeds motifs of these respective families into the same relative space, as shown in Figure 4, further illustrating the confounding nature of these motifs. A similar trend is found for motifs phosphorylated by PKC, PKA, and Akt. This trio is also shown to be closely related as indicated by the correlations in Figure 3 and the embeddings in Figure 4.

**Fig. 5.**
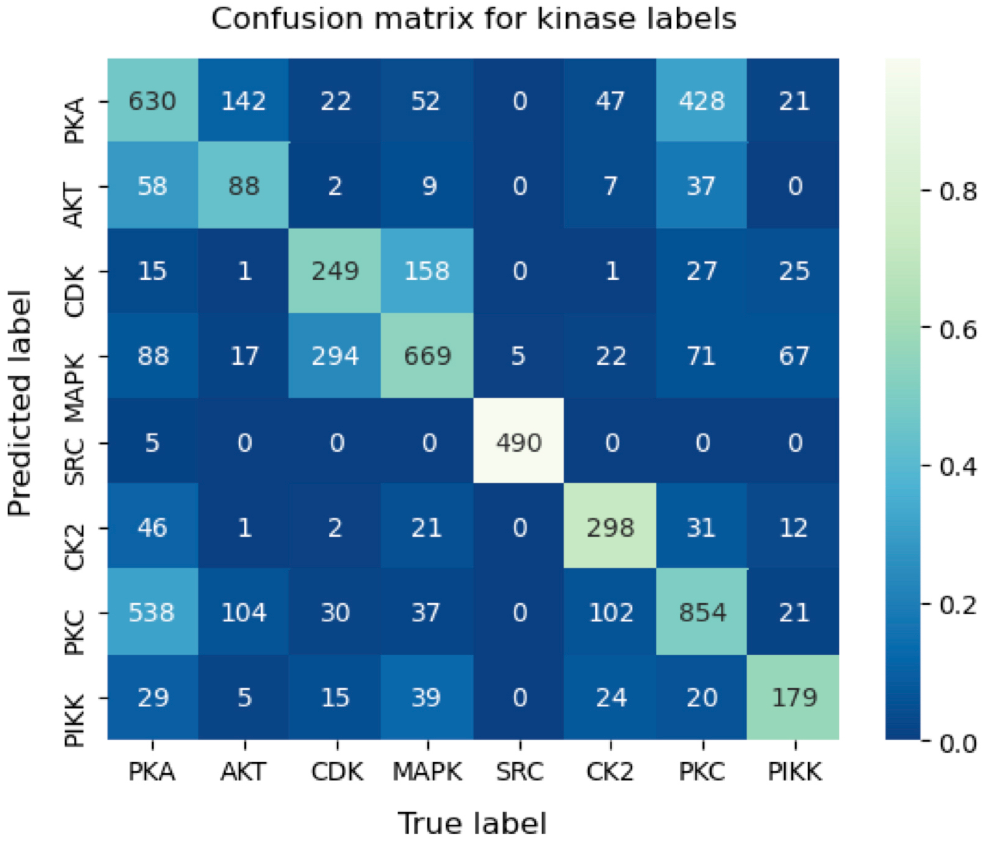
Confusion matrix for EMBER predictions on the test set. The numbers inside each box represent the raw number of predictions per box. The color scale is based on the ratio of predictions (in the corresponding box) to total predictions, per label. A lighter color corresponds to a larger ratio of predictions to total predictions.

**Fig. 6.**
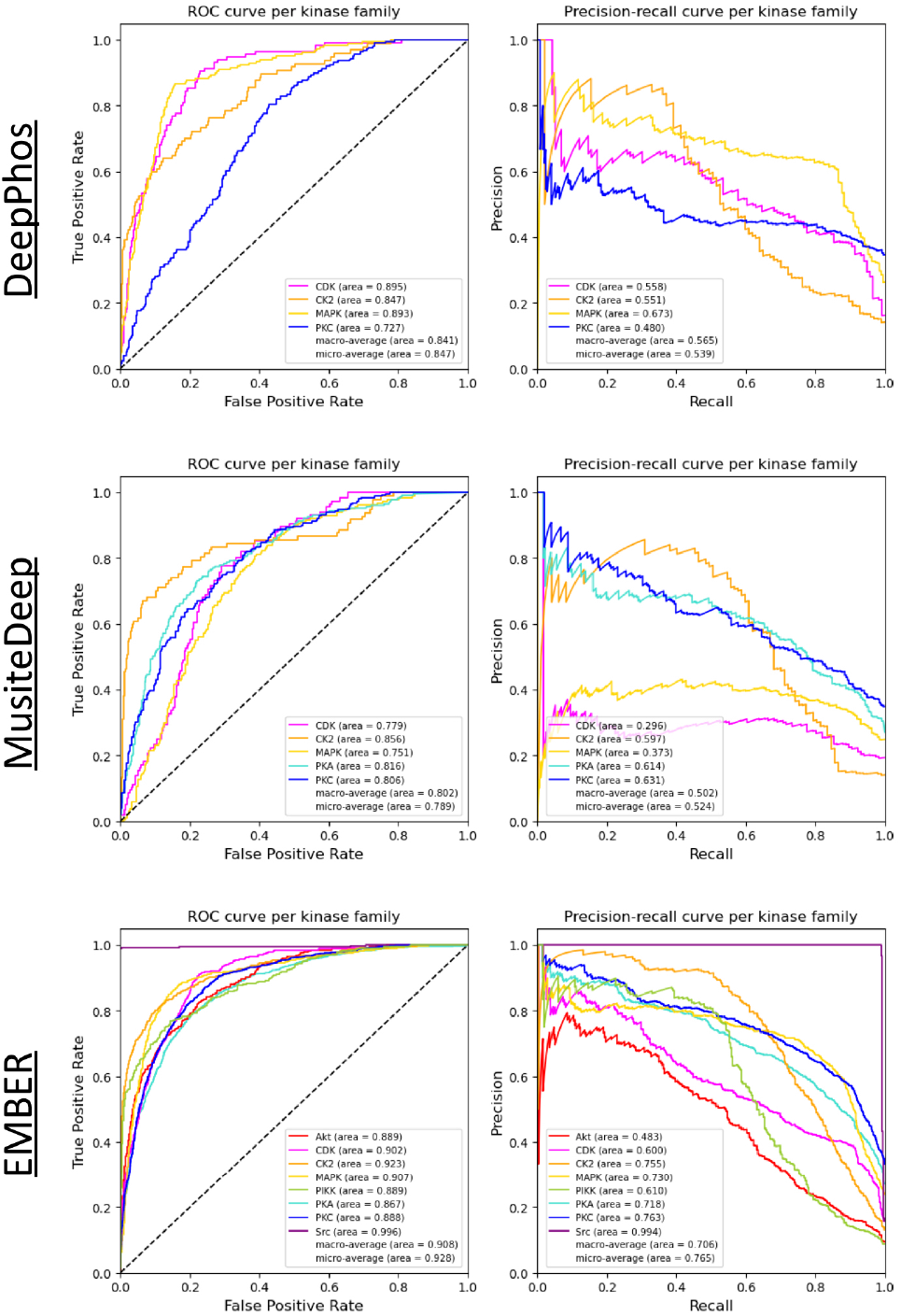
AUROC and AUPRC results achieved on the independent test set by DeepPhos, MusiteDeep, and EMBER. The AUROC and AUPRC of each kinase family label is shown in the respective legends.

#### Comparison to existing methods

We sought to compare EMBER’s performance to the two existing deep learning methods, MusiteDeep and DeepPhos, which adopt single-label models. However, this is not a straight-forward comparison because EMBER was trained on sequences 15 amino acids in length while MusiteDeep and DeepPhos were trained on sequences of 33 and 51 amino acids in length, respectively. Thus, we must elongate our 15-mers to lengths of 33 and 51 in order for MusiteDeep and DeepPhos to accept those sequences into their architectures as input. To accomplish this, we queried the Uniprot database to find complete protein sequences of which our test set motifs were subsequences. For instances in which a motif was a subsequence of multiple proteins we chose a protein at random from the set. By referencing the original complete protein sequence we were able to elongate our motifs by adding nine (in the case of MusiteDeep) or 18 (in the case of DeepPhos) amino acids to each flank of the original 15-mer motif. This resulted in a test set of 33-mers for MusiteDeep and 51-mers for DeepPhos.

We note that of the eight kinase families for which our model produces predictions, DeepPhos has functioning models for only four of the families (CDK, CK2, MAPK, and PKC), and MusiteDeep has models for only five of the families (CDK, CK2, MAPK, PKA, and PKC). We show AUROC and AUPRC results per kinase label from each of the three methods in Figure 6. EMBER outperforms MusiteDeep and DeepPhos on all four averaged metrics, indicating that our multi-label approach may be better equipped to solve the problem of kinase-motif prediction compared to the single-label approaches.

## Discussion

Illuminating the map of kinase-substrate interactions has the potential to enhance our understanding of basic cellular signaling as well as drive health applications, for example, by facilitating the development of novel kinase inhibitor-based therapies that disrupt kinase signaling pathways. Here, we have presented a deep learning-based approach that aims to predict which substrates are likely to be phosphorylated by a specific kinase family. In particular, our multi-label approach establishes a unified model that utilizes all available kinase-motif data to learn broader structures within the data and improve predictions across all kinase families in tandem. This approach avoids challenges in hyperparameter tuning inherent in the development of an individual model for each kinase. We believe that this approach will enable continuing improvement in predictions, as newly generated data describing any kinase-motif phosphorylation event can assist in improving predictions for all kinases. That is, a kinase-motif interaction discovered for PKA will improve the predictions not just for PKA, but also for Akt, PKC, MAPK, etc. through the transfer learning capabilities inherent in our multi-label model.

We showed that incorporation of a learned vector representation of motifs, namely the motifs’ coordinates in the Siamese embedding space, serves to improve performance over a model that utilizes only one-hot encoded motif sequences as input. Not only did the Siamese embedding improve prediction of phosphorylation events through a neu ral network architecture, but it also outperformed ProtVec, a previously developed embedding, in a coordinate-based *k*- NN task. This improvement over ProtVec was in spite of the fact that the Siamese network utilized less than 7,000 training sequences of 15 amino acids in length compared to ProtVec’s 500,000 sequences of approximately 300 amino acids in average length. The Siamese embedding was further generated through direct comparison of kinase-motif phosphorylation events rather than simply the sequence-derived data used by ProtVec. Furthermore, ProtVec is a generalized protein embedding while the Siamese embedding described here has the potential to be customized. For example, the use of the Jaccard distance in the Siamese loss allows the network to be trained on any number of multi-label datasets such acetylation, methylation, and carbonylation reactions. We also found that there is a small though meaningful relationship between the evolutionary distance between kinases and the motifs they phosphorylate, supporting the concept that closely related kinases will tend to phosphorylate similar motifs. When encoded in the form of our phylogenetic loss function, this relationship was able to slightly improve prediction accuracies. Together, these results suggest that EMBER holds significant promise towards the task of illuminating the currently unknown relationships between kinases and the substrates they act on.

## ACKNOWLEDGEMENTS

We would like to acknowledge members of the GomezLab for helpful comments and feedback. This work was supported by grants through the National Institutes of Health (Grant #s CA177993, CA233811, CA238475, DK116204).

## Supplementary Materials

### EMBER: Multi-label prediction of kinase-substrate phosphorylation events through deep learning

#### 1. Metrics

Definitions of metrics that characterize the area under the receiver operator curve (AUROC) and the precision-recall curve (AUPRC) as described in the main manuscript:

The AUROC is the integral of the receiver operator curve, which is found by plotting the true positive rate (TPR) against the false positive rate (FPR) at various decision thresholds. The TPR is defined as TP / (TP + FN) where TP are the true positive predictions and FN are the false negative predictions. The FPR is defined as FP / (FP + TN) where FP are the false positive predictions and TN are the true negative predictions.

The AUPRC is the integral of the receiver operator curve, which is found by plotting the precision against the recall (i.e. TPR) at various decision thresholds. Precision is defined as: TP / (TP + FP) where TP are the true positive predictions and FP are the false positive predictions.

#### 2. ProtVec embedding figures

Here, we show a qualitative comparison, via a UMAP reduction, between the original ProtVec embedding and the averaged ProtVec embedding.

##### Original

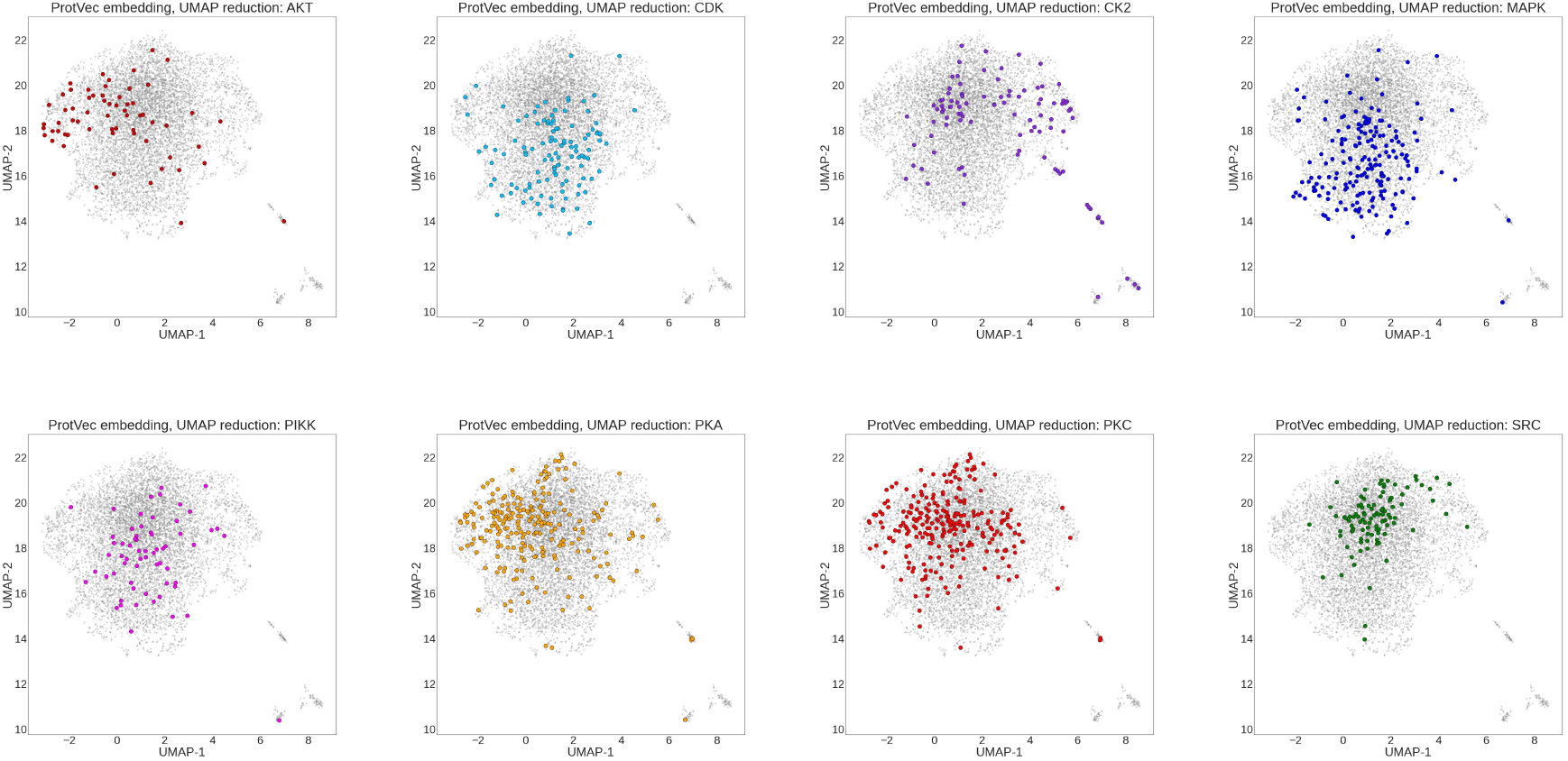

##### Averaged

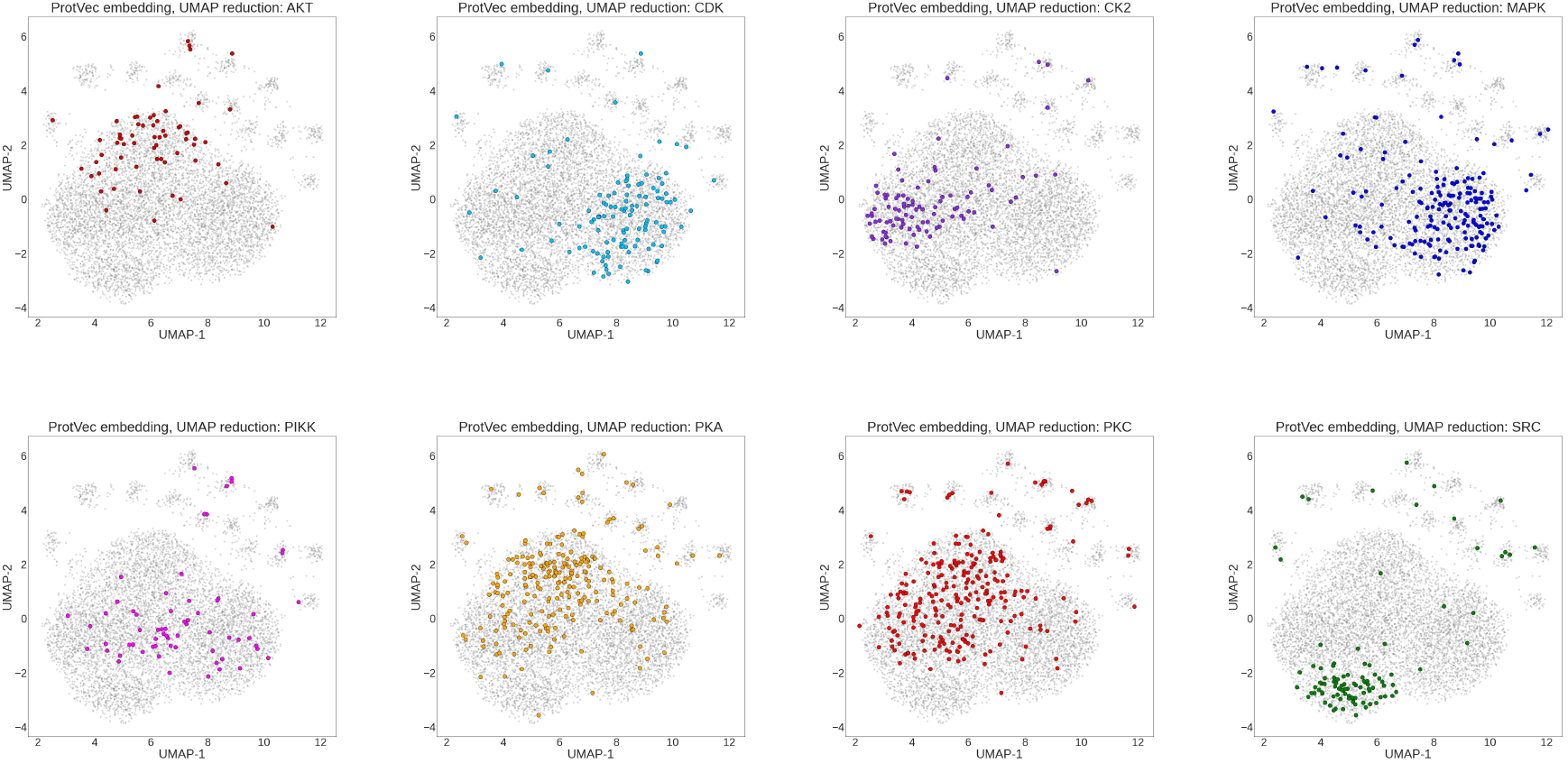

#### 3. ProtVec embedding kNN results

In the table below we show the AUROC and AUPRC results of the kNN classification task on the original ProtVec embedding and the averaged ProtVec embedding. For our kNN calculation, we used k = 85.

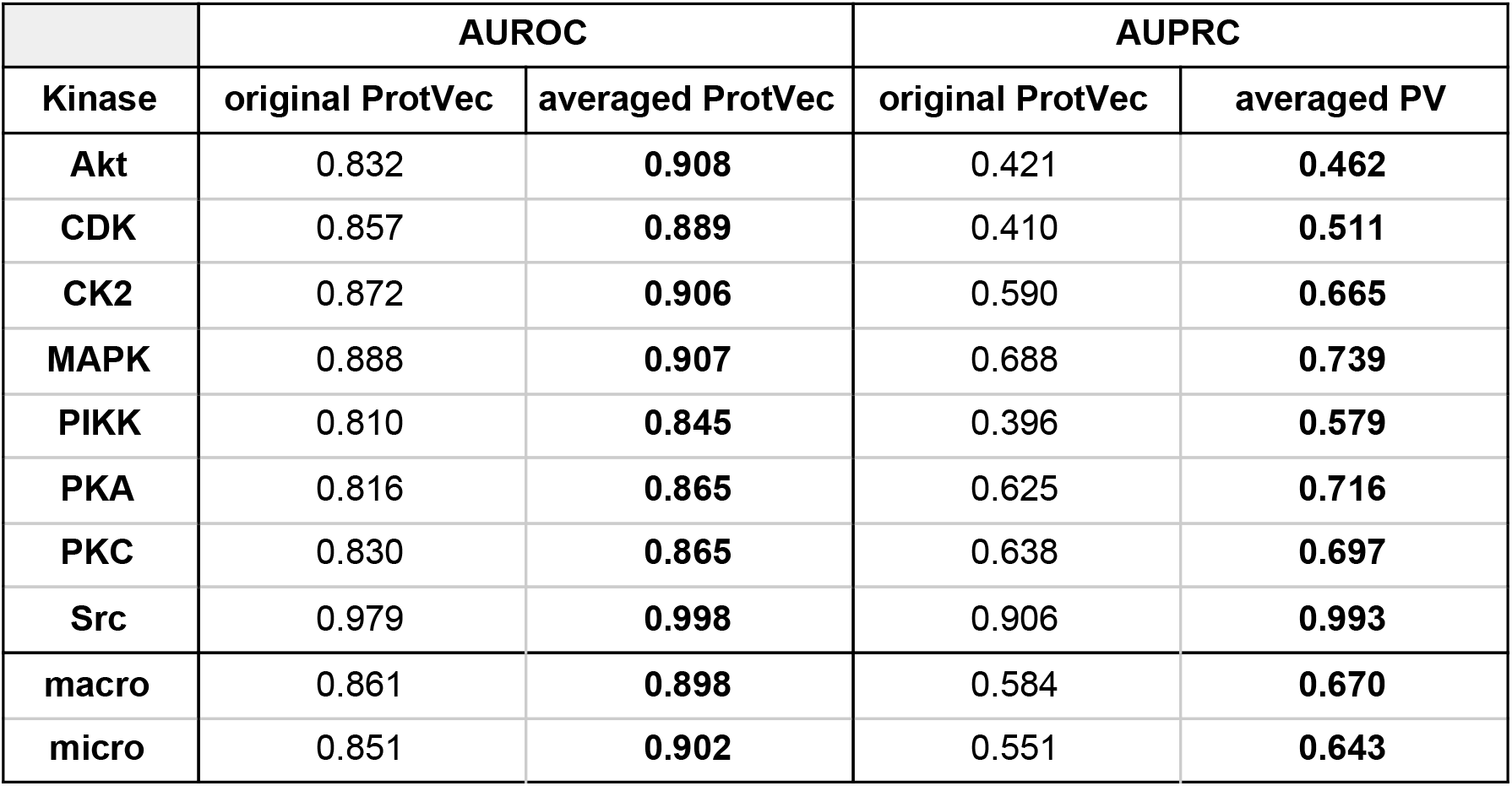

| Kinase | AUROC |  | AUPRC |  |
| --- | --- | --- | --- | --- |
|  | original ProtVec | averaged ProtVec | original ProtVec | averaged PV |
| **Akt** | 0.832 | **0.908** | 0.421 | **0.462** |
| **CDK** | 0.857 | **0.889** | 0.410 | **0.511** |
| **CK2** | 0.872 | **0.906** | 0.590 | **0.665** |
| **MAPK** | 0.888 | **0.907** | 0.688 | **0.739** |
| **PIKK** | 0.810 | **0.845** | 0.396 | **0.579** |
| **PKA** | 0.816 | **0.865** | 0.625 | **0.716** |
| **PKC** | 0.830 | **0.865** | 0.638 | **0.697** |
| **Src** | 0.979 | **0.998** | 0.906 | **0.993** |
| **macro** | 0.861 | **0.898** | 0.584 | **0.670** |
| **micro** | 0.851 | **0.902** | 0.551 | **0.643** |

#### 4. Hardware

Training and testing of EMBER occurred on a Linux system with the following configuration:

- Pop!_OS Linux 20.04
- Intel Xeon E5-2620 v4 with 32 cores @ 3.0 GHz
- 128 GB Ram
- Nvidia Titan Xp

On this system, the Siamese network took around 12 minutes to train, and the classification network took around 6 minutes to train.

## Bibliography

1. Tzong-Yi Lee, Justin Bo-Kai Hsu, Wen-Chi Chang, and Hsien-Da Huang. RegPhos: a system to explore the protein kinase-substrate phosphorylation network in humans. Nucleic Acids Res., 39(Database issue):D777–87, January 2011.

2. G Manning, D B Whyte, R Martinez, T Hunter, and S Sudarsanam. The protein kinase complement of the human genome. Science, 298(5600):1912–1934, December 2002.

3. Panayotis Vlastaridis, Pelagia Kyriakidou, Anargyros Chaliotis, Yves Van de Peer, Stephen G Oliver, and Grigoris D Amoutzias. Estimating the total number of phosphoproteins and phosphorylation sites in eukaryotic proteomes. Gigascience, 6(2):1–11, February 2017.

4. Leah J Wilson, Adam Linley, Dean E Hammond, Fiona E Hood, Judy M Coulson, David J MacEwan, Sarah J Ross, Joseph R Slupsky, Paul D Smith, Patrick A Eyers, and Ian A Prior. New perspectives, opportunities, and challenges in exploring the human protein kinome. Cancer Res., December 2017.

5. G L Johnson and Razvan Lapadat. Mitogen-Activated protein kinase pathways mediated by ERK, JNK, and p38 protein kinases. Science, 298(5600):1911, 2002.

6. Gayathri K Perera, Chrysanthi Ainali, Ekaterina Semenova, Christian Hundhausen, Guillermo Barinaga, Deepika Kassen, Andrew E Williams, Muddassar M Mirza, Mercedesz Balazs, Xiaoting Wang, Robert Sanchez Rodriguez, Andrej Alendar, Jonathan Barker,Sophia Tsoka, Wenjun Ouyang, and Frank O Nestle. Integrative biology approach identifies cytokine targeting strategies for psoriasis. Sci. Transl. Med., 6(223):223ra22, February 2014.

7. Nicole Tegtmeyer, Matthias Neddermann, Carmen Isabell Asche, and Steffen Backert. Subversion of host kinases: a key network in cellular signaling hijacked by helicobacter pylori CagA. Mol. Microbiol., May 2017.

8. Amandine Charras, Pinelopi Arvaniti, Christelle Le Dantec, Marina I Arleevskaya, Kaliopi Zachou, George N Dalekos, Anne Bordon, and Yves Renaudineau. JAK inhibitors suppress innate epigenetic reprogramming: a promise for patients with sjögren’s syndrome. Clin. Rev. Allergy Immunol., June 2019.

9. Alessia Alunno, Ivan Padjen, Antonis Fanouriakis, and Dimitrios T Boumpas. Pathogenic and therapeutic relevance of JAK/STAT signaling in systemic lupus erythematosus: Integration of distinct inflammatory pathways and the prospect of their inhibition with an oral agent. Cells, 8(8), August 2019.

10. Ya Nan Deng, Joseph A Bellanti, and Song Guo Zheng. Essential kinases and transcriptional regulators and their roles in autoimmunity. Biomolecules, 9(4), April 2019.

11. Kyla A L Collins, Timothy J Stuhlmiller, Jon S Zawistowski, Michael P East, Trang T Pham, Claire R Hall, Daniel R Goulet, Samantha M Bevill, Steven P Angus, Sara H Velarde, Noah Sciaky, Tudor I Oprea, Lee M Graves, Gary L Johnson, and Shawn M Gomez. Proteomic analysis defines kinase taxonomies specific for subtypes of breast cancer. Oncotarget, 9 (21):15480–15497, March 2018.

12. Elise J Needham, Benjamin L Parker, Timur Burykin, David E James, and Sean J Humphrey. Illuminating the dark phosphoproteome. Sci. Signal., 12(565), January 2019.

13. Wenwen Fan, Xiaoyi Xu, Yi Shen, Huanqing Feng, Ao Li, and Minghui Wang. Prediction of protein kinase-specific phosphorylation sites in hierarchical structure using functional information and random forest. Amino Acids, 46(4):1069–1078, April 2014.

14. Shu-Yun Huang, Shao-Ping Shi, Jian-Ding Qiu, and Ming-Chu Liu. Using support vector machines to identify protein phosphorylation sites in viruses. J. Mol. Graph. Model., 56: 84–90, March 2015.

15. Fuyi Li, Chen Li, Tatiana T Marquez-Lago, André Leier, Tatsuya Akutsu, Anthony W Purcell, A Ian Smith, Trevor Lithgow, Roger J Daly, Jiangning Song, and Kuo-Chen Chou. Quokka: a comprehensive tool for rapid and accurate prediction of kinase family-specific phosphorylation sites in the human proteome. Bioinformatics, 34(24):4223–4231, December 2018.

16. Yu Xue, Ao Li, Lirong Wang, Huanqing Feng, and Xuebiao Yao. PPSP: prediction of PK-specific phosphorylation site with bayesian decision theory. BMC Bioinformatics, 7:163, March 2006.

17. Duolin Wang, Shuai Zeng, Chunhui Xu, Wangren Qiu, Yanchun Liang, Trupti Joshi, and Dong Xu. MusiteDeep: a deep-learning framework for general and kinase-specific phosphorylation site prediction. Bioinformatics, 33(24):3909–3916, December 2017.

18. Fenglin Luo, Minghui Wang, Yu Liu, Xing-Ming Zhao, and Ao Li. DeepPhos: prediction of protein phosphorylation sites with deep learning. Bioinformatics, 35(16):2766–2773, August 2019.

19. Ehsaneddin Asgari and Mohammad R K Mofrad. Continuous distributed representation of biological sequences for deep proteomics and genomics. PLoS One, 10(11):e0141287, November 2015.

20. Peter V Hornbeck, Jon M Kornhauser, Sasha Tkachev, Bin Zhang, Elzbieta Skrzypek, Beth Murray, Vaughan Latham, and Michael Sullivan. PhosphoSitePlus: a comprehensive resource for investigating the structure and function of experimentally determined post-translational modifications in man and mouse. Nucleic Acids Res., 40(Database issue): D261–70, January 2012.

21. Jianfei Hu, Hee-Sool Rho, Robert H Newman, Jin Zhang, Heng Zhu, and Jiang Qian. PhosphoNetworks: a database for human phosphorylation networks. Bioinformatics, 30(1):141–142, January 2014.

22. Holger Dinkel, Claudia Chica, Allegra Via, Cathryn M Gould, Lars J Jensen, Toby J Gibson, and Francesca Diella. Phospho.ELM: a database of phosphorylation sites–update 2011. Nucleic Acids Res., 39(Database issue):D261–7, January 2011.

23. Jane Bromley, Isabelle Guyon, Yann LeCun, Eduard Säckinger, and Roopak Shah. Signature verification using a “siamese” time delay neural network. In J D Cowan, G Tesauro, and J Alspector, editors, Advances in Neural Information Processing Systems 6, pages 737–744. Morgan-Kaufmann, 1994.

24. R Hadsell, S Chopra, and Y LeCun. Dimensionality reduction by learning an invariant mapping. In 2006 IEEE Computer Society Conference on Computer Vision and Pattern Recognition (CVPR’06), volume 2, pages 1735–1742, June 2006.

25. Leland McInnes, John Healy, and James Melville. UMAP: Uniform manifold approximation and projection for dimension reduction. February 2018.

